# Conditioning by Subthreshold Synaptic Input Changes the Intrinsic Firing Pattern of CA3 Hippocampal Neurons

**DOI:** 10.1101/084152

**Authors:** Saray Soldado-Magraner, Federico Brandalise, Suraj Honnuraiah, Michael Pfeiffer, Urs Gerber, Rodney Douglas

## Abstract

Unlike synaptic strength, intrinsic excitability is assumed to be a stable property of neurons. For example, learning of somatic conductances is generally not incorporated into computational models, and the discharge pattern of neurons in response to test stimuli is frequently used as a basis for phenotypic classification. However, it is increasingly evident that signal processing properties of neurons are more generally plastic on the timescale of minutes. Here we demonstrate that the intrinsic firing patterns of CA3 neurons of the rat hippocampus in vitro undergo rapid long-term plasticity in response to a few minutes of only subthreshold synaptic conditioning. This plasticity on the spike-timing could also be induced by intrasomatic injection of subthreshold depolarizing pulses and was blocked by kinase inhibitors, indicating that discharge dynamics are modulated locally. Cluster analysis of firing patterns before and after conditioning revealed systematic transitions towards adapting and intrinsic burst behaviours, irrespective of the patterns initially exhibited by the cells. We used a conductance-based model to decide appropriate pharmacological blockade, and found that the observed transitions are likely due to recruitment of calcium and M-type potassium conductances. We conclude that CA3 neurons adapt their conductance profile to the subthreshold activity of their input, so that their intrinsic firing pattern is not a static signature, but rather a reflection of their history of subthreshold activity. In this way, recurrent output from CA3 neurons may collectively shape the temporal dynamics of their embedding circuits.

**New & Noteworthy:** Despite being widely conserved across the animal phyla, it is still a mystery why nerve cells present diverse discharge dynamics upon somatic step currents. Adding a new timing dimension to the intrinsic plasticity literature, here we show that CA3 neurons rapidly adapt through the space of known firing patterns in response to the subthreshold signals that they receive from their embedding circuit. This result implies that CA3 neurons collectively adjust their network processing to the temporal statistics of their circuit.

## Introduction

Neurons of the Central Nervous System present an intriguing diversity of firing patterns in response to applied intra-somatic step currents (McCormick et al., 1985; Connors and Gutnick, 1990; Cauli et al., 2000; Markram et al., 2004; Butt et al., 2005; Dumitriu et al., 2007; Hemond et al., 2008; Somogyi and Klausberger, 2005; Tasic et al., 2016). These responses may be for example: adapting, accelerating, bursting or fast spiking, but why neurons have evolved these diversity of precise spike-timings remains an open issue. With some exceptions (Steriade, 2004), the patterns are assumed to be a sufficiently stable property of a neuron to be used as a basis for phenotypic classification (Markram et al., 2004; Ascoli et al., 2008; Tricoire et al., 2011; Van Aerde and Feldmeyer, 2015).

However, there are substantial reasons to doubt that firing patterns are static properties of neurons. The discharge dynamics depends on the distribution and activations of the membrane conductances that it expresses (Hille, 2001; Markram et al., 2004). This distribution is subject, over the course of hours and days, to homeostatic control via up- or down-regulation of conductances in response to the neuron’s own activity (Turrigiano et al., 1994; Turrigiano and Nelson, 2004; Marder and Goaillard, 2006). Furthermore, neurons have conserved molecular pathways that link network activity to the recruitment of genes and signaling factors implicated in neural excitability (Flavell and Greenberg, 2008; Cohen and Greenberg, 2008), and activity-dependent maturation is necessary for the emergence of the whole spectrum of electrical types (Moody and Bosma, 2005; García et al., 2011). These lines of evidence suggest that the firing pattern is not a static characteristic of the cell, but rather the consequence of adaptive mechanisms that adjust the spike-timing of the neuron in response to the patterns of activity in its embedding network. In fact, some studies show that specific features of the discharge response are actually modulated homeostatically upon activity or after learning. For example, the spike delay (Cudmore et al., 2010; Dehorter et al., 2015) or the accommodation rate (Thompson et al., 1996).

Changes in intrinsic excitability have recently been reported also on faster time scales (Aizenman and Linden, 2000; Paz et al., 2009; Mahon and Charpier, 2012; Brager and Johnston, 2007). These shifts are postulated to increase the probability of firing, integrating neurons into memory engrams in synergy with hebbian learning (Zhang and Linden, 2003; Titley et al., 2017). However, the fact that key conductances with potential impact on the spike pattern have also been found to be rapidly modulated by activity (Belmeguenai et al., 2010; Hyun et al., 2013; Brown and Randall, 2009; Grasselli et al., 2016) suggests neurons may be doing more than just increasing their overall excitability, for instance, adjusting their temporal processing as well (Debanne et al., 2019). Despite the evidence, it remains unexplored whether neurons can rapidly modulate upon activity not just specific features of the discharge or its average rate, but the whole spectrum of intrinsic spike-timings classically described (Ascoli et al., 2008).

We have addressed this issue by studying the effect that subthreshold activity has on the suprathreshold response of neurons using whole-cell recordings in the CA3 region of the rat hippocampus in organotypic cultures. The discharge patterns were characterized before and after a conditioning phase of periodic subthreshold synaptic stimulation lasting a few minutes. Pre-conditioned cells presented diverse discharge patterns (Ascoli et al., 2008). However, conditioning by subthreshold synaptic input elicited significant long-lasting changes in the behavior of most of the neurons examined, requiring substantial re-classification of their type. This effect was reproduced when conditioning the cells via subthreshold intra-somatic current pulses and was blocked by adding protein kinase A (PKA) and protein kinase C (PKC) inhibitors to the recording pipette, suggesting that changes are mediated at the single cell level via phosphorylation. The conditioning effect was also reproduced in juvenile mice using the acute slice preparation technique, indicating it is a generic property of CA3 cells. Using a conductance-based neuron model and the channel blockers *XE*991 and *NiCl*_2_ we found that the results can be explained by a recruitment of voltage dependent calcium and M-type potassium conductances. We conclude that CA3 neurons rapidly adapt their spike-timing response by tuning their conductance profile to the subthreshold inputs of their embedding circuit.

## Results

### Firing patterns of CA3 neurons change after subthreshold synaptic stimulation

Whole-cell patch clamp recordings of CA3 neurons were performed in rat hippocampal organotypic cultures. The intrinsic firing patterns of the neurons were recorded before and after conditioning by extracellular stimulation of the mossy fibers (MF) originating in the dentate gyrus. The conditioning stimuli consisted of paired pulses (0.1 ms duration pulses, with an interval of 10 – 20 ms) applied at 1 Hz, and repeated 500 times for a total period of approximately 8 minutes. The amplitude of the pulses was adjusted for each recorded cell to elicit only subthreshold excitatory post-synaptic potentials (EPSPs). This mossy fiber stimulation protocol is a modification of that described by Brandalise and Gerber (2014) and Brandalise et al. (2016), which has been previously shown to elicit heterosynaptic subthreshold plasticity in CA3 pyramidal-pyramidal synapses. We tested whether the same type of subthreshold stimulation could also induce plasticity on the action potential discharge patterns of the CA3 cells. The firing patterns of neurons were assessed with a sequence of constant current injections. For convenience, we label these patterns according to the Petilla classification terminology (Ascoli et al., 2008). Interestingly, we observed that for most neurons the conditioning protocol elicited a change in the Petilla discharge pattern, independently of their type of firing on control conditions. For example, the pyramidal cell shown in Figure 1A had a non-adapting burst pattern before stimulation (grey traces) but after conditioning (blue traces), this response changed to intrinsic burst. The same transition was observed for the pyramidal cell on Figure 1B, whose initial pattern was delayed accelerating. The bipolar cell on Figure 1C switched from non-adapting continuous to adapting continuous firing. The most common transition was towards adapting and intrinsic burst patterns. This observation is supported by measuring the mean fraction of spikes in the first half versus the second half of the voltage traces for the population of recorded cells. The distribution of the spikes favors the first half (Figures 1D-E) (n=50), supporting our observation that the main pattern transitions are towards adapting and intrinsic burst behaviors after the conditioning.

**Figure 1.**
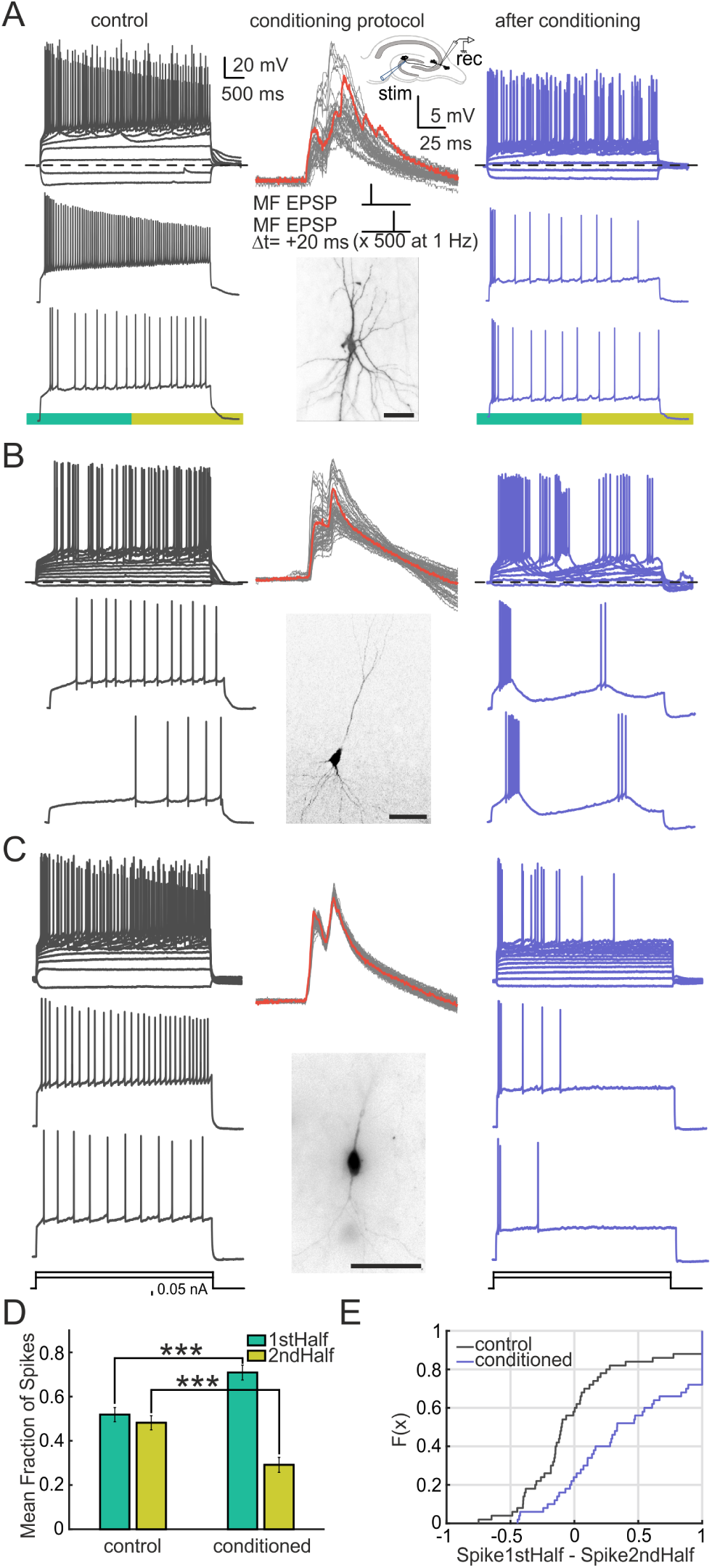
Firing pattern transitions occur in CA3 neurons after subthreshold paired-pulse stimulation of afferents. A-C) Three examples of neurons in the CA3 area presenting different morphologies and different firing patterns in control conditions. The discharge patterns were measured by injection of step currents of increasing amplitude. Control measurements (grey traces, left) were followed by stimulation of the mossy fibers. The upper trace shows all voltage traces elicited upon different levels of current injection on that cell. Two sample traces of this set are shown below. EPSPs (middle panel) were evoked in response to a stimulation with double current pulses, separated by 20 ms and repeated 500 times at 1 Hz. The series of repeated pulses are shown superimposed. The median trace is highlighted in red. The inset shows the configuration of recording and stimulating electrodes (on the CA3 region of the hippocampus and on the dentate gyrus, respectively). Below, the morphology obtained by labelling the cells with biocytin is shown. After the conditioning, patterns were measured again (blue traces, right). A) Pyramidal cell switches from non-adapting burst to intrinsic burst firing. B) Pyramidal cell switches from delay accelerating to intrinsic burst continuous pattern. C) Bipolar cell switches from non-adapting continuous to adapting continuous firing (scale bars = 50*µ*m). D) Mean fraction of spikes for the population in the first and second half of the voltage trace (see green and yellow rectangle below the trace in A for an example) for both control and conditioned cases. A significant redistribution on the fraction of spikes is observed after the conditioning, where the fraction of spikes on the first half is increased while it decreases in the second half (n=50, p=1.92e-6, two-sided Wilcoxon signed rank test). E) Empirical cumulative distribution function for the data shown in D. Every individual cell, for both control and conditioned cases, is represented as the number of spikes for the first half of the trace minus the spikes for the second half (n=50)

The mossy fiber conditioning was followed by a significant 36 MΩ (25%) decrease in input resistance (Rin), (from 144.8 ± 73.0MΩ to 108.4 ± 65.3MΩ, two-sided Wilcoxon signed rank test, p=1.1e-5), while there was no significant change in rheobase (from 0.36 ± 0.32 nA to 0.3 ± 0.6 nA, two-sided Wilcoxon signed rank test, p=0.59). There was also a significant 5 mV (7%) depolarization of the resting membrane potential (Vm) (−65.3 ± 5.0mV) with respect to resting level (−70.4 ± 5.7mV, two-sided Wilcoxon signed rank test, p=2.3e-5, n = 50). However, the firing pattern transitions could not be induced by simply clamping the membrane potential at different values (see Figure S1D and F, n = 10), nor could they be induced by the step-currents used to measure the discharge patterns (see Figure S1A-C, n = 15). No significant changes in Vm or Rin were found in unconditioned cells (Vm: -69.3 ± 2.0mV, -69.1 ± 1.9mV, paired t-test, p=0.64, Rin: 148.8 ± 56.1MΩ, 158.9 ± 55.6MΩ, paired t-test, p=0.063, n = 15). Intracellular dialysis could also be excluded as the cause of the pattern transitions, as firings did not change spontaneously over intracellular recording time (see Figures S1D and F). In addition, a change in the fraction of spikes in favor of the first half was also found after conditioning in a setting where dialysis was minimized (high resistance pipette recordings, 10-12 MΩ, n=10, p=0.048, two-sided Wilcoxon signed rank test).

We noted however a significant change in the mean fraction of spikes (in favor of the first half) after 20 min of recording, when assessing the firing pattern three times during this period (n = 12, p=0.005, two-sided Wilcoxon signed rank test). This indicates that transient suprathreshold current steps can also trigger changes in firing, as previous studies in CA3 have reported (Brown and Randall, 2009), although continuous subthreshold conditioning over a shorter time scale has a stronger effect (8 minutes, Figure 1).

### Firing pattern transitions are gradual during the course of the conditioning and long-lasting

We next assessed whether the expression of the firing transition was gradual or whether firings changed more sharply during the course of the conditioning. To this end, we repeated the original experiment (Figure 1) while assessing the firing pattern of the cells every 100 trials of conditioning. The general observation was that cells displayed a gradual transition in firing, in which more regular or accelerating patterns changed towards different degrees of adaptation and intrinsic burst responses. For example, the cell shown in Figure 2A presents a non-adapting or regular pattern in control conditions. After 100 repetitions of the conditioning protocol the spike distribution changes to an adapting continuous pattern. This adaptation gets reinforced after 200 conditionings, with the cell later adopting an intrinsic burst pattern. Other cells showed a progression for more regular to only adapting patterns. As an average, the smooth transition is reflected in the mean fraction of spikes shown in Figures 2B and C. As it can be observed, successive conditionings translate into a stronger redistribution of this fraction in favor of the first half of the trace, a reflection of the cells adopting a stronger adapting or intrinsic burst responses (n=8). The progression of the resting membrane potential (Vm) and input resistance (Rin) during the course of the conditioning was also monitored, and it is shown in Figure 2D. A progressive increase in Vm and a progressive decrease in Rin is observed (Rin, from 166.4 ± 54.2MΩ to 115.3 ± 53.3MΩ, two-sided Wilcoxon signed rank test, p=0.031, n=7) (Vm, from -64.6 ± 6.2mV to -58.7 ± 5.7mV, two-sided Wilcoxon signed rank test, p=0.047, n = 7).

**Figure 2.**
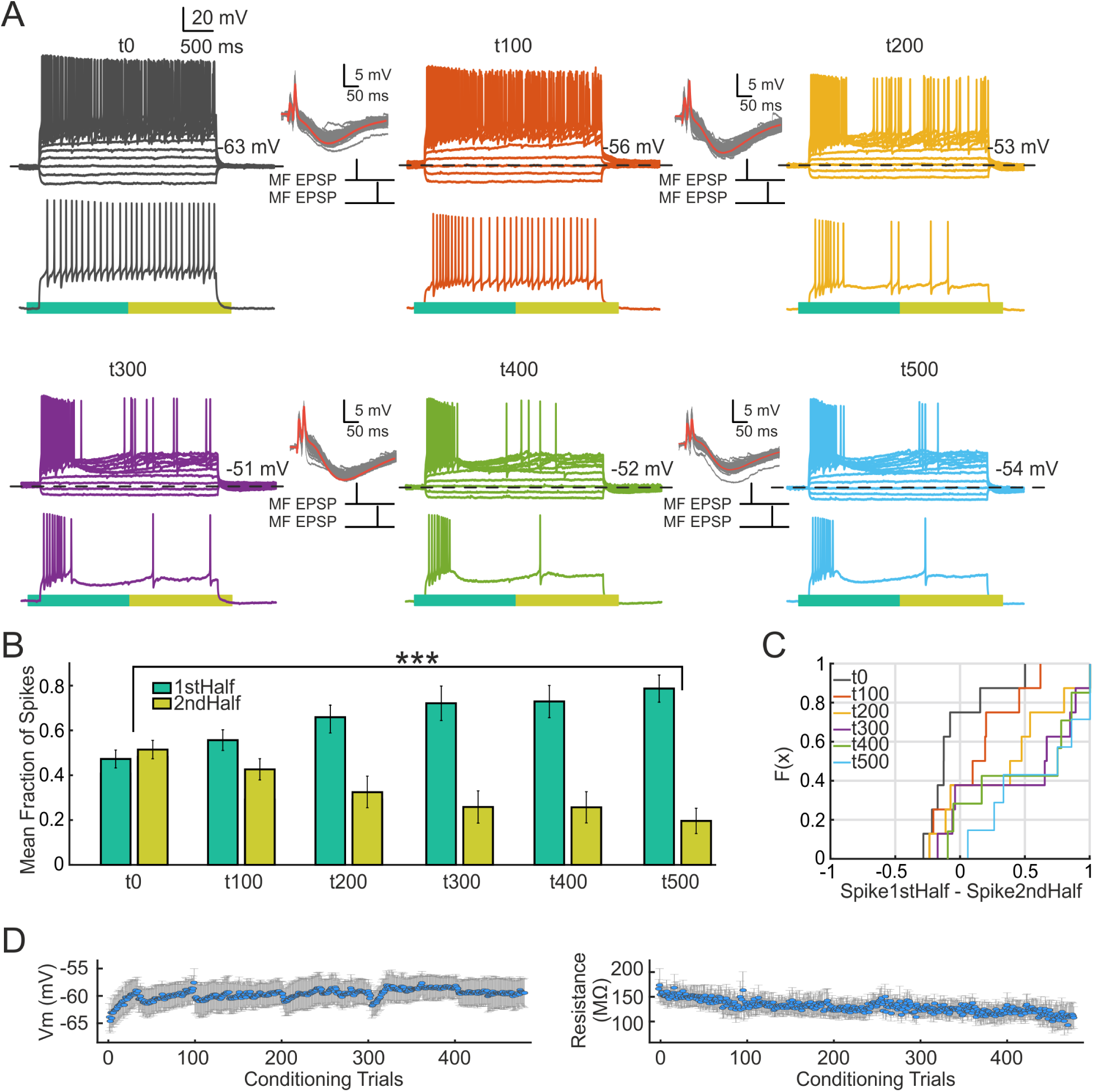
The expression of the conditioning effect is gradual over the course of the stimulation. Firing patterns were assessed every 100 conditionings until 500 trials were completed. A) Representative cell whose firing pattern is non-adapting in control conditions (grey). After 100 stimulations the cell shows an adapting pattern (red), after 200 the adaptation gets stronger and an intrinsic burst pattern emerges after the successive conditionings (orange to blue). Conditioning protocols are showed on the insets. Red line shows the median. B) Mean fraction of spikes for the population in the first and second half of the voltage trace during the successive conditionings. A significant redistribution on the fraction of spikes is observed during the course of the conditioning. The fraction of spikes on the first half continuously increases in favor of the second half (n = 8, p=1.24e-5, repeated measures ANOVA) C) Empirical cumulative distribution function for the data shown in B. The number of spikes for the first half of the trace minus the spikes for the second half is shown for every cell (n=8) D) Average resting membrane potential (Vm) and input resistance (Rin) of the cells during the course of the conditioning. Vm was measured as the baseline voltage before the depolarization caused by the conditioning, Rin was measured by injecting a negative current step after each conditioning trial. Circles indicate mean, bars indicate SEM (n=8). In the cases where Vm was above -60 mV a small holding current was injected to keep the cell stable during the conditioning.

Changes in intrinsic properties of neurons have been previously reported to be long-lasting, similar to synaptic forms of long-term plasticity (Turrigiano and Nelson, 2004; Titley et al., 2017). We thus assessed whether the firing pattern plasticity was stable in time or whether the changes were transient. We conditioned the cells and followed their firing response up to 30 minutes after conditioning. In all cases, cells presented a post-conditioned characteristic change in firing pattern towards adapting and intrinsic burst patterns (as in Figure 1). This same pattern was assessed every 10 minutes and showed to be persisting up to the whole recording period (see Figure 3), indicating a long-lasting change in intrinsic excitability. In two cells the pattern persisted up to 50 min of recording. A representative cell is shown in Figure 3A, which presented an accelerating pattern that switched to intrinsic burst after conditioning. This induced pattern remained stable for 30 min. At the population level, the stability is reflected in the mean fraction of spikes. After the conditioning effect, no significant change in this fraction was observed over 30 min (see Figure 3C)(n=6). Note that for this subset of cells a 60 ms paired pulse was employed during the conditioning (see Figure 3A). The previous observed changes in firing occurred upon conditioning with a paired mossy fiber current pulse separated by 20 ms and given at a frequency of 1 Hz (Figure 1). We wondered whether a different timing interval (60 ms) would affect the outcome of the conditioning, as shown by previous reports of subthreshold synaptic plasticity in the CA3 circuit (Brandalise and Gerber, 2014; Brandalise et al., 2016). However, no differential effect on the pattern was found (Figure 3), indicating that the mechanism of intrinsic firing plasticity is not sensitive to these two different time scales.

**Figure 3.**
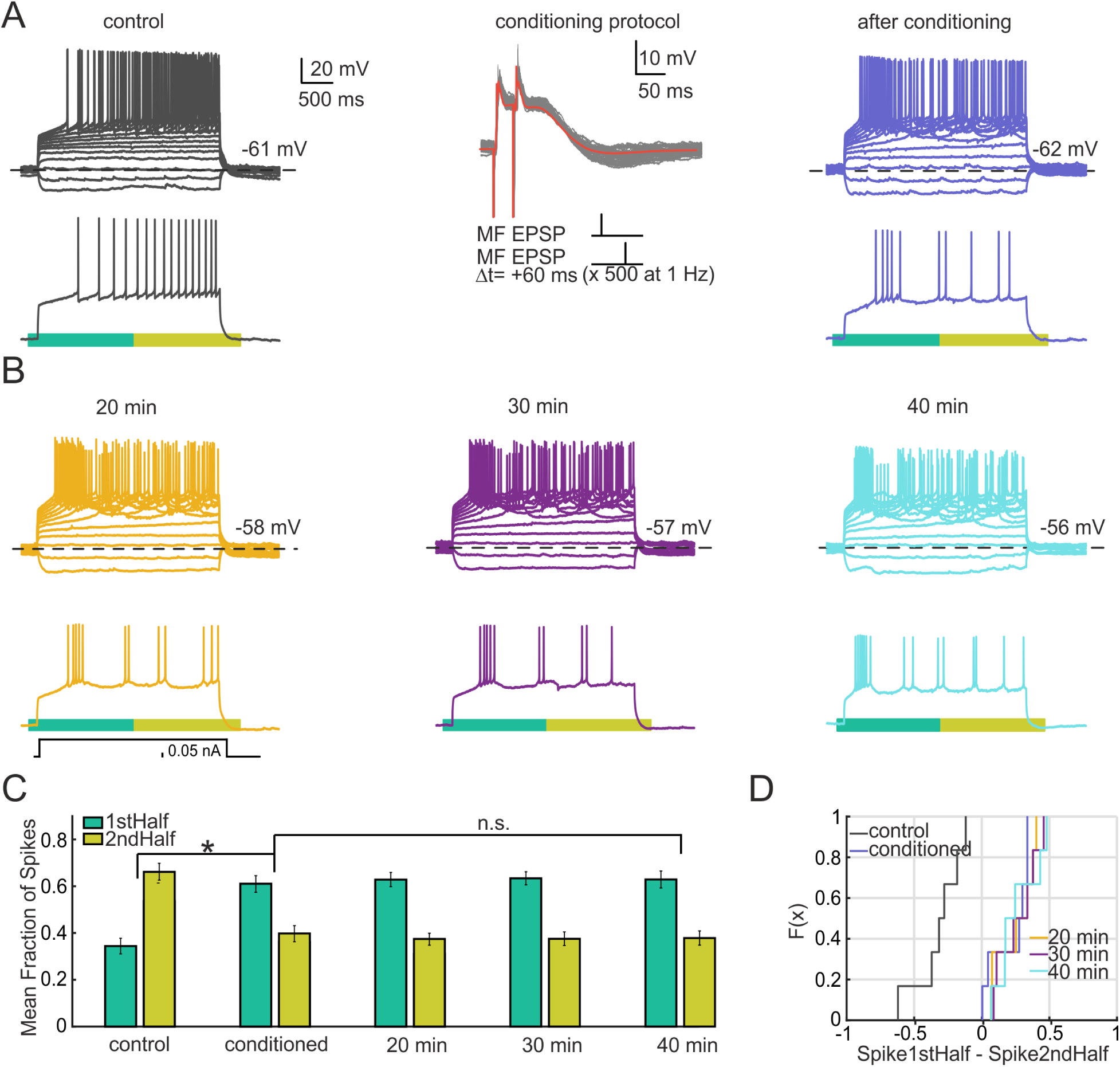
The effect of the conditioning is persistent over time. Cells where followed for 40 minutes to assess whether the firing pattern plasticity was long-term. A) Example cell with an accelerating firing pattern in control conditions (grey). The cell was conditioned subthresholdy with a double mossy fiber current pulse separated by 60 ms and given at a frequency of 1 Hz (protocol is shown in the middle, red line indicates the median). A change in pattern to intrinsic burst is elicited (blue). B) The same cell was followed every 10 min after conditioning until reaching 40 min of recording (orange, purple and blue). The pattern remained stable. C) Mean fraction of spikes for the population in the first and second half of the voltage trace before, after conditioning and every 10 min thereafter. A significant redistribution on the fraction of spikes is observed after the conditioning (n=6, p=0.031, two-sided Wilcoxon signed rank test). No significant change in this fraction was observed over 30 min after conditioning (n=6, p=0.4, repeated measures ANOVA). D) Empirical cumulative distribution function for the data shown in C.

### Firing pattern transitions are independent of synaptic input and are blocked by protein kinase A and C inhibitors

We attempted to resolve whether synaptic input was necessary to elicit the changes, or whether they could be induced by direct stimulation of the soma. To this end, we used intra-somatic injection of paired step current pulses whose parameters were chosen to elicit a similar somatic voltage response compared to that generated by the mossy fiber stimulation (Figure 4). This direct subthreshold somatic stimulus evoked changes in discharge pattern that were similar to those elicited by the indirect mossy stimulation. For example, the cell in Figure 4A displayed a delay accelerating firing pattern in control conditions and underwent a transition towards intrinsic burst pattern after somatic conditioning. The population data for direct stimulation showed a significant redistribution in the fraction of spikes in favor of the first half of the trace versus the second half after the conditioning (Figures 4B and C) (n=12). In these experiments we observed the same tendency of neurons to become adapting and intrinsic burst after conditioning as for mossy fiber stimulation. This result suggests that the mechanism underlying the changes in firing pattern is not localized to synapses, but rather acts at a more central, probably somatic or proximal dendritic level.

**Figure 4.**
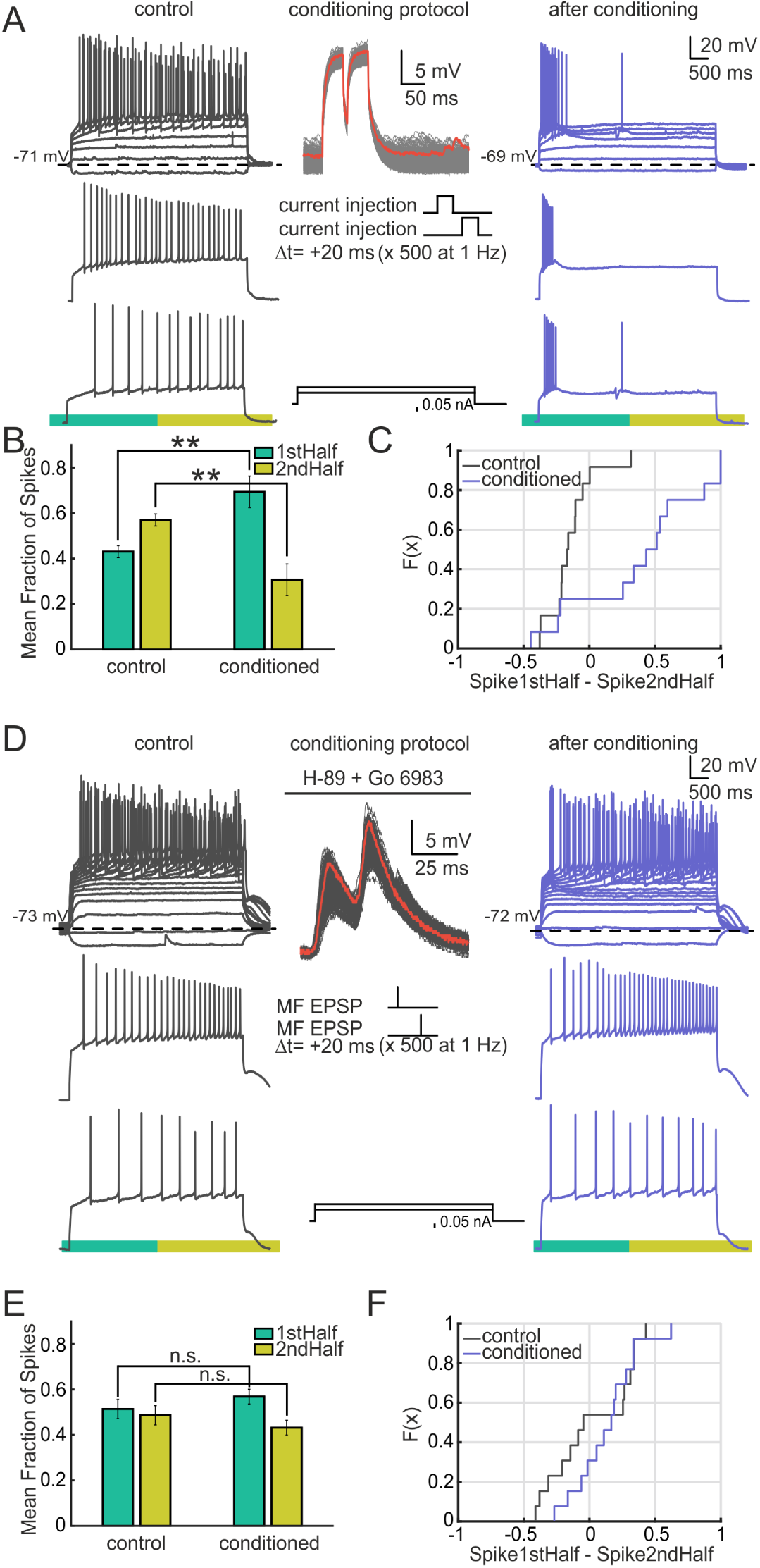
CA3 firing pattern transitions occur upon somatic conditioning and are blocked by kinase inhibitors. A) Example of an intrasomatic conditioned cell that switched from delay accelerating (grey traces) to intrinsic burst firing (blue traces). The conditioning protocol is shown in the middle column. The red line shows the median. EPSPs were evoked by injection of paired current steps, of 50 ms in duration and separated by 20 ms. The double steps were repeated 500 times at 1 Hz. The series of repeated pulses are shown superimposed. A sample trace is shown in red. B) Mean fraction of spikes for the population in the first and second half of the voltage trace for both control and conditioned cases. A significant redistribution on the fraction of spikes occurs after the conditioning. The fraction of spikes on the first half is increased while it decreases in the second half (n=12, p=0.0024, two-sided Wilcoxon signed rank test). C) Empirical cumulative distribution Function for the data shown in B. Every individual cell is represented as the number of spikes for the first half of the trace minus the spikes for the second half (n=12). D) Example of a mossy fiber conditioned cell (as described in Figure 1) under the presence of H-89 and Go 6983 (PKA and PKC inhibitors) on the intracellular pipette. The cell expresses a delay accelerating pattern in control conditions and remains under such pattern after the conditioning protocol is applied. E) Mean fraction of spikes for the population in the first and second half of the voltage trace for both control and conditioned cases. The redistribution of the fraction of spikes was not significant after the conditioning (n=13, p=0.266, two-sided Wilcoxon signed rank test). F) Empirical cumulative distribution function for the data shown in D. Every individual cell is represented as the number of spikes for the first half of the trace minus the spikes for the second half (n=13).

The fact that the firing pattern transitions could be reproduced by this direct depolarization of the soma raised the question of whether the somatic depolarization elicited by mossy fiber activation is necessary to elicit the observed changes in firing. We thus repeated the mossy conditioning experiment (Figure 1) while artificially hyperpolarizing the neuron (see Figure S2). The hyperpolarization did not abolish the effect of the conditioning, since a significant redistribution of spikes in favor of the first half of the trace was observed after the conditioning (see Figure S2B and C). However, the effect was stronger when conditioning the cells without hyperpolarization (see Figure S2B or Figure 1D for comparison). An example cell is shown in Figure S2A. The cell presents an accelerating pattern in control conditions. After conditioning via the mossy fiber pathway, under the presence of an hyperpolarizing pulse, the cell changed its firing towards a non-adapting burst pattern. When re-conditioning the cell with no hyperpolarization the pattern switched to intrinsic burst. These results suggest that a transient depolarization, such as the intrasomatically injected stimulus, is sufficient but not necessary to elicit the effect. The residual effect may indicate that the mechanism is localized near the MF synapse, in which case somatic hyperpolarization could be insufficient to prevent depolarization there. Note however that in most neurons a handful of depolarizing trials were accidentally elicited while adjusting the magnitude of the current MF pulse. Additionally, some cells presented occasional rebound spiking caused by the hyperpolarization, while an increase in stimulation amplitude due to the increase in driving force was also frequent. This could all potentially contribute to the observed effect.

Next, we sought to identify what internal mechanism could be responsible for the firing pattern transitions. The firing pattern of the cell depends on the distribution of ion channels, or conductances, that the cell presents at its membrane (Hille, 2001). A possible mechanism would act by changing this distribution. Due to the time scale of the response (on the order of minutes) we ruled out protein synthesis of new channels on the membrane. An alternative could be channel phosphorylation, a mechanism known to affect the maximal conductance of the channel on a relatively short timescale (Davis et al., 2001). We reproduced the conditioning protocol in the presence of the PKA and PKC inhibitors H-89 and Go 6983 in the intracellular recording pipette. Figure 4D shows a cell whose firing pattern in control conditions was delay accelerating. After mossy fiber conditioning in the presence of the inhibitors the cell retained this initial pattern. 84% of cells recorded with phosphorylation inhibitors showed no visible modulation of the Petilla label pattern (11 out of the 13 cells). Figures 4E and F show the population response for these cells. Unlike Figure 1D, no significant redistribution of the spikes was found on the presence of the inhibitors (n=13). These results suggest that phosphorylation is implicated in the mechanism of firing pattern transition.

### Using Dynamic Time Warping (DTW) and a conductance based model to infer firing transitions and parameter changes after plasticity

We observed that the conditioning induced firing pattern changes from more regular patterns towards early bursting and adapting patterns. We sought to quantify these changes using hierarchical clustering methods (Druckmann et al., 2013; Tricoire et al., 2011; Hosp et al., 2014) to establish more objectively which discharge type to associate with every response, and to quantify the frequencies of transitions between them. For our clustering method, we obtained instantaneous firing rate vectors of the experimental voltage traces and estimated pairwise distances using the DTW algorithm. DTW operates directly on the action potential temporal sequence rather than relying on a pre-defined set of features (Druckmann et al., 2013; Tricoire et al., 2011; Hosp et al., 2014). The rate vectors used for the clustering can be interpreted as the subthreshold voltage envelope in which the discharge response of each cell rides, and this envelope is the essence to catalogue similar firing patterns. Once the distances between the vectors were calculated, Wards linkage was applied in order to obtain a hierarchical tree to reveal the classes.

The results of the cluster analysis of discharge patterns are shown in Figure 5A. We set the threshold of the clustering tree at a level that separates the traces into 5 distinct families. The threshold was chosen large enough to yield sufficient structure to interpret the hierarchy in terms of recognized response types (Ascoli et al., 2008). Representative traces of each family are shown in Figure 5B. The average of the firing rate vectors of every cluster is depicted beneath each representative trace.

**Figure 5.**
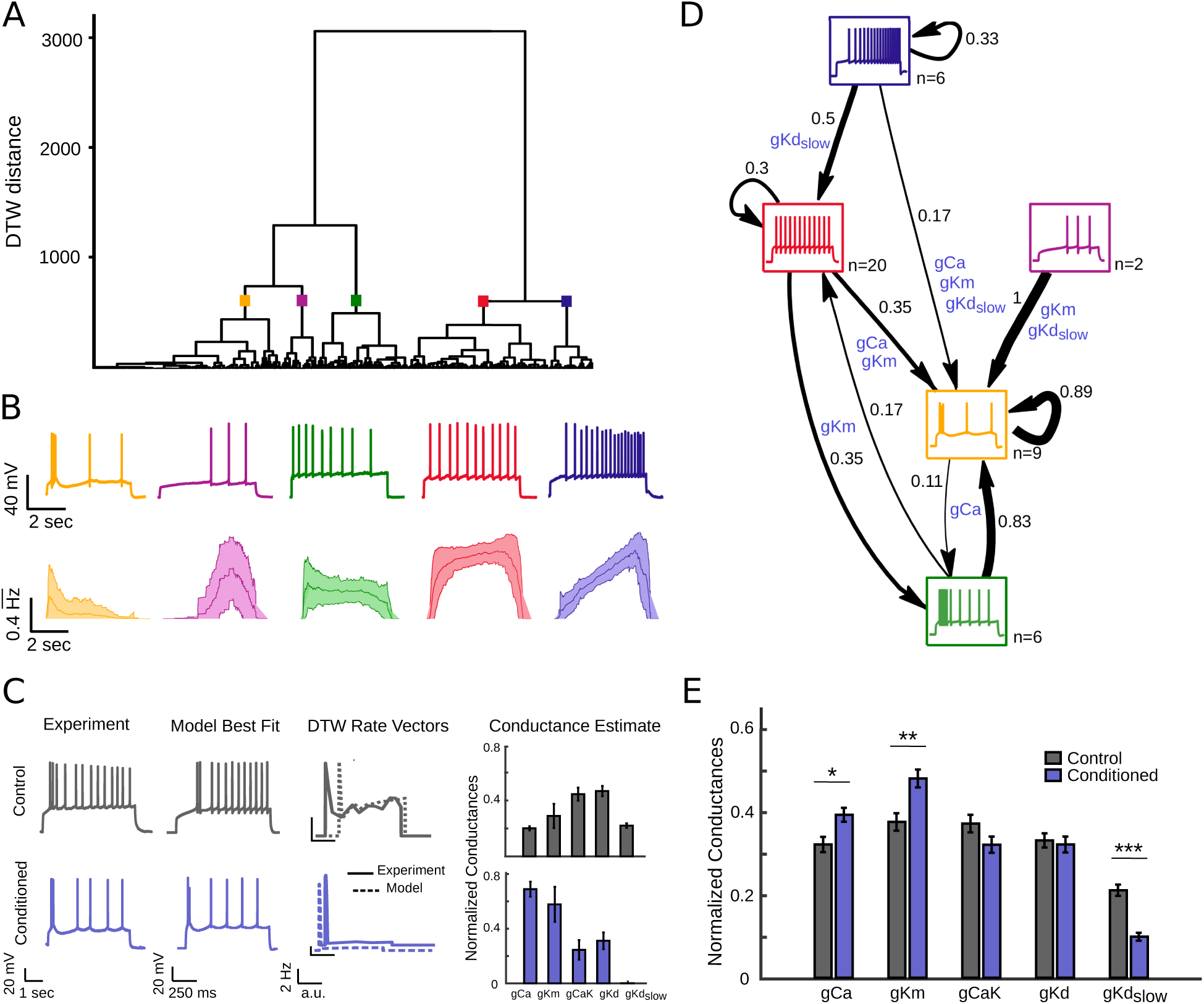
Hierarchical clustering of experimental discharge traces and mapping to a conductance model. A) Dendrogram of clustered traces. The data included in the cluster corresponds to the mossy fiber conditioned cells of Figure 1. Two main families can be identified: one containing adapting and bursting traces, together with delayed spiking patterns (left branch); and another branch containing regular and accelerating traces (right branch) (n=50). B) Representative traces from each cluster. Below, average instantaneous firing rate over all traces belonging to the same cluster. Middle lines indicate the mean; light outer lines indicate standard deviations. The instantaneous firing rate (in Hz) is normalized to 1. C) Every experimental trace is matched to a model database of traces. Using the DTW distance on the instantaneous firing rate vectors the best matches are selected (best match is depicted). A conductance estimate for the experimental trace is obtained (average of 10 best matches are shown). D) Transitions observed between firing patterns before and after conditioning. Each cell is assigned to a single cluster (represented as a box) for both the control and conditioned cases. Arrows indicate transitions between types whenever a cell changed cluster. Self-loops indicate that the firing pattern was retained after conditioning. Numbers indicate percentages of observed transitions. The number of cells in each category under control conditions is displayed next to each pattern type. Cells tend to transition towards adapting and bursting patterns following conditioning (n = 43). Seven cells were assigned as unclassified. A conductance road map showing the key conductances responsible for a transition in firing pattern are represented on the edges. The main channels implicated are *gCa*, *gKd_slow_* and *gKm*. E) Average conductance composition for matched experimental cells in control (grey) and conditioned cases (blue). There is a significant increase in *gKm*, *gCa* and a decrease in *gKd_slow_*(*gCa* p=0.015, *gKm* p= 0.0084, *gCaK* p= 0.2203, *gKd* p= 0.2501, *gKd_slow_* p=2.01*e^−^*^8^, two-sided Wilcoxon signed rank test, n = 485).

The clustering algorithm successfully captures the typical time courses of the firing patterns. The right branch of the cluster tree contains accelerating and non-adapting firing patterns, while the other contains adapting and intrinsic bursting patterns together with a smaller group of traces that have delayed spiking profiles. The consistency of the algorithm was confirmed by its successful clustering of independent rate vectors derived from the same set of current injections (same cell under the same conditions) into a single cluster. Indeed, in 86% of cases (43 of the 50 cells) the algorithm successfully allocated the majority of vectors from the same set of current injections into single clusters. Vectors from the 7 remaining cells were not consistently classified. For 50% of the cells all of their voltage traces fell into the same cluster, and for 90% of the cells at least 50% did. The allocation of some responses from the same cell into more than a single cluster does however follow a biological logic. For example, for cells classified as accelerating, some of their voltage traces could reasonably fall into the non-adapting cluster because acceleration may vanish at high current injections. A similar reasonable misclassification is possible for adapting traces. In this case low current injections may be classified as non-adapting because the currents are not high enough to elicit adaptation. In particular, many of the traces belonging to the delayed spiking cluster are derived from cells whose traces at low current injections were assigned to the accelerating cluster, or belonged to non-adapting cells with spiking delay. The transitions between cluster types induced by the stimulation protocol are shown in Figure 5D. This figure considers only those cells in which responses both before and after conditioning could be clearly assigned to a cluster. In total, 68% of the cells (n = 50) changed their original cluster as a result of subthreshold conditioning. This quantitative result supports the observation that cells tend to transition towards more adapting and intrinsic burst profiles. 70% of cells initially belonging to the non-adapting cluster exhibited such changes in response (14 cells), with 35% moving into the intrinsic burst category, and 35% exhibiting adapting spike patterns. 5 of the 6 cells from the adapting cluster (83%) switched to the intrinsic burst type. Most of the cells for which the firing pattern did not change were already in the most common target states of transitions. For example, 89% of the intrinsic bursting cells did not change cluster. This provides further evidence for a predominantly unidirectional change of firing patterns in response to conditioning. The 7 cells that could not be consistently classified under control conditions were all correctly classified after the stimulation. They showed the same transition tendencies: 5 moved into the intrinsic bursting cluster, the other 2 became adapting.

We next aimed to infer which underlying parameters could be responsible for the systematic transitions. Our results showing that phosphorylation inhibition blocks the conditioning effect support the hypothesis that the prime candidate for this plasticity is a change in the profile of active conductances. We explored this possibility using simulations of action potential discharge in a conductance-based single compartment neuron model containing 10 voltage and calcium gated ion channels (see Methods). The densities and kinetics of these channels were derived from experimental measurements of CA3 pyramidal neurons (Hemond et al., 2008).

A database of representative ranges of maximal conductances that could plausibly explain the discharge patterns observed experimentally was generated using the single compartment model. To do this, the maximal conductances of the different channels were swept through ranges that would likely encompass the experimentally observed patterns. The spiking conductances were left constant, whereas we varied the conductances with longer time constant, which are responsible of the discharge dynamics: *gCa*, *gCaK*, *gKm*, *gKd* and *gKd_slow_* (see Table 1 for the exact ranges). In this manner a total of 100’000 conductance profiles were generated. We obtained the discharge response to different levels of current injection for each conductance profile, giving a total of 800’000 voltage traces with their associated maximal conductance profiles. Every single experimental trace (coming from both, control and conditioned cases) was matched against the collection of traces in the model database using the DTW algorithm on the instantaneous firing rate vectors (see Figure 5C for an example). The best fits were then selected, allowing us to obtain an estimate of the maximal conductance profile likely to be present in the experimental neuron (Figure 5C). The key to infer the parameters is thus to recognize, via the DTW algorithm on the rate vectors, the subthreshold voltage envelope generated by the long-time constant conductances, and not the precise spike times.

**Table 1.** Range of maximal conductance values used to generate the model database of voltage traces. A model database of voltage traces, which includes all the observed experimental firing patterns, was generated by varying 5 maximal conductances (*gCa*, *gCaK*, *gKm*, *gKd* and *gKd_slow_*) over a given range. Different ranges of step current *I* were also needed to reveal the different firing types. A total of 100’000 conductance vectors were generated by combining the different conductances. The firing pattern of every conductance vector was produced at several levels of step-current injection, obtaining a total of 800’000 voltage traces. Note that *gCaT*, *gCaN* and *gCaL* are englobed under the single parameter *gCa*.

| Conductances and Current | Employed ranges |
| --- | --- |
| $g_{Na}$ | 0.04 |
| $g_{Kdr}$ | 0.01 |
| $g_{Ka}$ | 0.05 |
| $g_{Ca}$ | $1e^{-6} + (0 : 1e^{-4} : 9e^{-4})$ |
| $g_{CaK}$ | $1e^{-6} + (0 : 1e^{-4} : 9e^{-4})$ |
| $g_{Km}$ | $1e^{-6} + (0 : 1e^{-4} : 9e^{-4})$ |
| $g_{Kd}$ | $1e^{-6} + (0 : 1e^{-4} : 9e^{-4})$ |
| $g_{Kd_{slow}}$ | $1e^{-6} + (0 : 1e^{-4} : 9e^{-4})$ |
| $I$ | 0.2 : 0.16 : 1.32 |
| Total conductance vectors | 100'000 |
| Total traces | 800'000 |

The diagram of Figure 5D represents the crucial conductances determining the transitions between discharge patterns in firing pattern space. These are *gKm*, *gCa* and *gKd_slow_*. For example, to move to the intrinsic burst cluster (yellow) a characteristic enrichment in *gKm* and *gCa* is needed, which allows for the generation of the burst (given the presence of basal levels of *gCaK*) and the spacing of further spikes. For the accelerating and delayed patterns (blue and purple), the presence of *gKd* is important for a delayed onset of the spiking, and the slow inactivation of *gKd_slow_* is necessary for generating the continuous acceleration of the spike rate. In the case of the adapting patterns (green), the inclusion of *gKm* is necessary for the slowing down of the action potentials after the initial discharge.

Figure 5E shows the average conductance content of the matched experimental traces in control and conditioned cases. The shift towards adapting and intrinsic bursting behaviors after the conditioning corresponds to a significant increase in *gKm* and *gCa*, and a decrease in *gKd_slow_* (*gCa* p=0.015, *gKm* p= 0.0084, *gCaK* p= 0.2203, *gKd* p= 0.2501, *gKd_slow_* p=2.01*e^−^*^8^, two-sided Wilcoxon signed rank test, n = 485). This correspondence of firing patterns and biophysical parameters offers an interpretation of the causes of transitions between firing behaviours induced by the conditioning.

### Inhibition of Kv7 and calcium channels abolishes the effect of firing pattern plasticity

Having unraveled via the modeling study that the changes in firing pattern could correspond to an increase in *gKm* and *gCa* conductances, we decided to test this prediction via pharmacological blocking of the corresponding channels. We thus repeated the original experiment (Figure 1) and administrated the blockers *NiCl*_2_ and *XE*991 via the perfusion system after the conditioning. *NiCl*_2_ is known to block T,N and L type calcium channels, with a stronger effect on the T channel (Kochegarov, 2003), and *XE*991 is a selective blocker of the muscarinic potassium channel *Kv*7*/KCNQ* (Brown and Randall, 2009). The results of this experiment are shown in Figure 6. Administration of the drugs blocked the effect of the conditioning, with the cells loosing the change in pattern 20 min after perfusion (note that the drug took 3-4 min to reach the bath and start diffusing). For example, Figure 6A shows a cell that switched from accelerating to intrinsic burst after conditioning. Drugs were administrated immediately after checking the conditioning effect and the cell was followed for 30 min since that point (Figure 6C). 20 minutes after starting of drug perfusion the cell presented an adapting burst pattern, which became adapting continuous after 30 min. The population data shows a significant redistribution of spikes following drug perfusion, which goes in the opposite direction of that caused by the effect (see Figures 6E and F). Note that in the absence of the drugs, we have demonstrated that the plasticity is long-lasting over the course of these 30 min (Figure 3). Below each trace (Figures 6B and D), the model estimate of its conductance distribution is shown, as explained in Figure 5C. An increase in *gCa* and *gKm* conductances is observed after the conditioning, which then decreases after application of their corresponding channel blockers.

**Figure 6.**
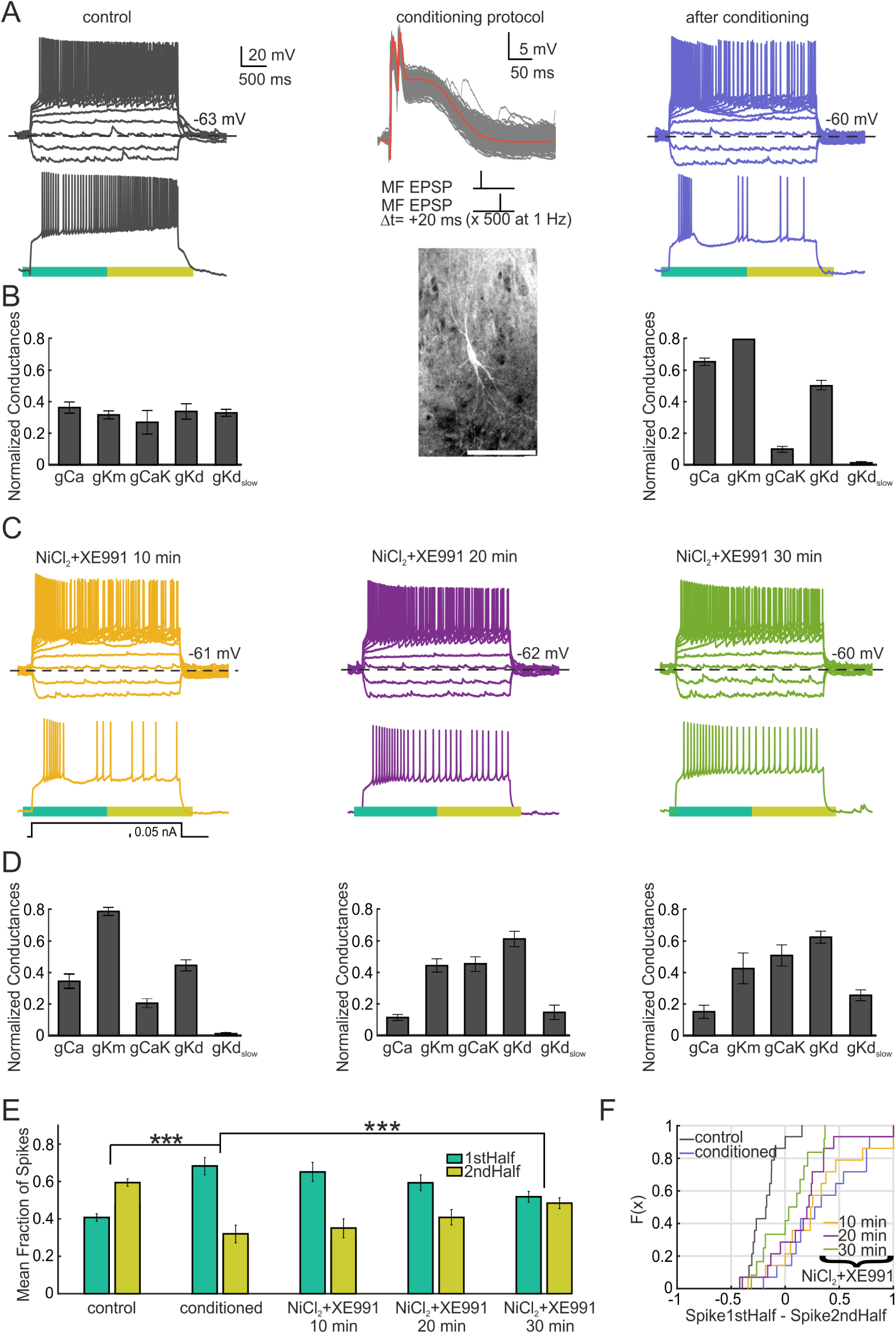
Inhibition of *Kv*7 and calcium channels abolish the effect of the conditioning. A) Example of a CA3 cell with accelerating pattern (grey). After conditioning via mossy fiber stimulation (protocol shown in the inset, red line shows the median.) the cell becomes intrinsic burst (blue). A biocytin staining of the cell is depicted below the protocol (scale bar = 200 *µ*m). C) After the conditioning, *NiCl*_2_ and *XE*991 were administered via the perfusion system (200 and 10 *µM*, respectively). The pattern was checked every 10 min since starting of perfusion. The drug took 3-4 minutes to reach the bath. 20 min since perfusion the cell presents an adapting burst pattern (purple). 10 min later the pattern can be catalogued as adapting continuous. B) and D) conductance distribution of every trace estimated via the model as done in Figure 5. Conditioning and consequent drug application affect the distribution of the conductance values. E) Mean fraction of spikes for the population in the first and second half of the voltage trace during the course of the experiment. A significant redistribution on the fraction of spikes towards the first half is observed after conditioning (n=14, p=0.0001, two-sided Wilcoxon signed rank test). This tendency is reverted during the drug application (n=14, p=7.2e-8, repeated measures ANOVA) F) Empirical cumulative distribution function for the data shown in E.

We noted that a residual adaptation remained in some neurons after the drug administration. This could likely be due to the low concentration of *XE*991 employed (10 *µm*), although it could also be due to the small acceleration at the beginning of the trace given by the delay present in these neurons. We decided to identify whether *gKd* was responsible for such delay as hinted by the model (Figure 6D). D-type currents, caused by Kv1 channels, can be blocked by low concentrations of 4-aminopyridine (4-AP). We thus repeated the experiment using 4-AP and *NiCl*_2_ as conductance blockers (see Figure S3). After successful conditioning of the cells (see Figure S3A for an example), the drugs were administered via the perfusion system. As in the previous experiment, the burst response presented by the neuron, likely caused by calcium channels, was abolished. 15 min after perfusion the delay presented by the neuron was also removed, with the cell adopting an adapting continuous pattern. Middle panel of Figure S3B shows the effect of 4-AP in removing the delay of the population of cells. There is a significant reduction of the timing of the first spike comparing to the *XE*991 experiment. For this subset of cells, no effect on the fraction of spikes was found after drug perfusion (see Figure S3E and F). This is likely due to the cells keeping adaptation profiles with no delay, in comparison with the previous set of experiments. Figures S3B and D show the corresponding conductance fits obtained from the voltage traces. The perfusion of the drugs results in a reduction of *gCa* and *gKd*, but not *gKm* as in the previous set of experiments. The present data indicate that the effect of firing pattern plasticity is likely being mediated by a recruitment of *gKm* and *gCa* conductances, with *gKd* necessary for shaping the delay of the spike response observed in the majority of the traces.

### Conditioning elicits a change in firing pattern on CA3 neurons in the acute slice preparation

This study was performed on organotypic cultures, derived from brain slices of newborn rats that were incubated for three weeks using the roller-tube technique (Gähwiler, 1981). Organotypic cultures have been used extensively to characterize electrophysiological properties of hippocampal neurons and it is known that the tissue preserves the anatomical organization of the adult hippocampus, as well as its connectivity and characteristic spontaneous activity (Gähwiler, 1988; Okamoto et al., 2014). However, hippocampal slices are derived from immature brains and this raises the question of whether the observed transitions of firing patterns are likely to happen in the mature tissue or whether they reflect activity-dependent acquisition of firing properties of neurons in the developing and more plastic brain. We thus decided to test whether this type of plasticity is also present on a different preparation, acute slices derived from mice between post natal days 15 and 22. Results are shown in Figure 7. Cells in the CA3 area were recorded and conditioned via somatic current injection (as in Figure 4). A subset of cells were also condioned via the mossy fiber pathway. In general, the conditioning elicited an increase in burstiness at the end of the voltage trace, which was higher in frequency of that encountered in the organotypic case. For example, the cell in Figure 7A presented a non-adapting pattern in control conditions. After conditioning the cell showed a delay burst pattern. A subset of cells also presented an increase in adaptation (Figure 7B) as observed in the organotypic slice case (Figure 5D). A significant change in the fraction of spikes was found at the population level (Figures 7C and D). These results indicate that conditioning elicits a change in firing pattern in the acute slice preparation, suggesting the firing pattern plasticity is a generic property of CA3 cells.

**Figure 7.**
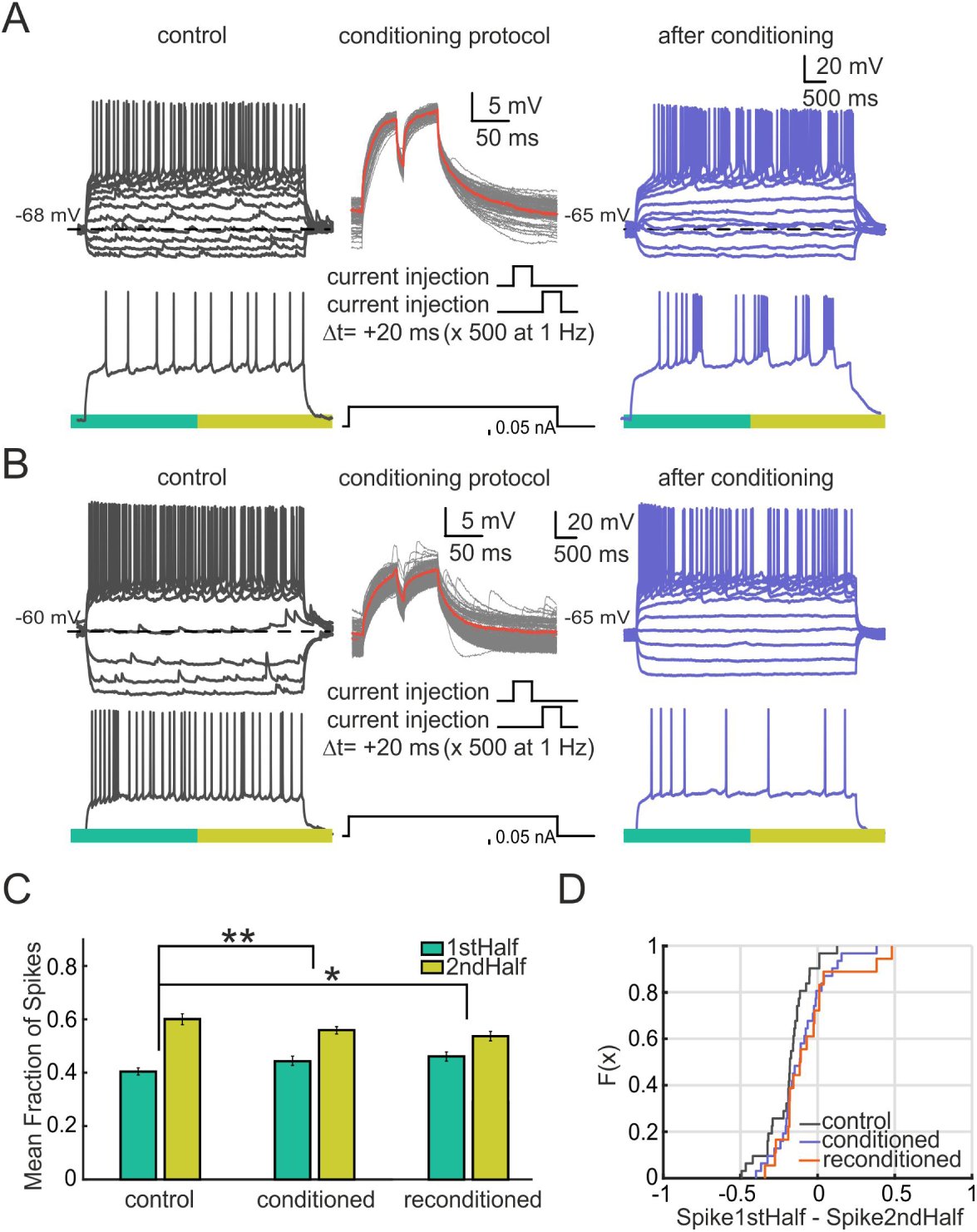
Conditioning changes the firing pattern of CA3 neurons in the acute slice preparation. A) Example of a cell with a non-adapting firing pattern in control conditions (grey). The cell is conditioned via somatic current injections (as in Figure 4). Protocol is shown in the inset. The firing pattern of the cell changes towards delay bursting (blue). B) Cell presenting a non-adapting burst firing pattern in control conditions. After somatic conditioning the cell presents an adapting firing response. C) Mean fraction of spikes for the population in the first and second half of the voltage trace during successive conditionings. There is a significant redistribution of the fraction of spikes after the conditioning (control-conditioned, n = 31, p=0.003, two-sided Wilcoxon signed rank test) (control-conditioned-reconditioned, n = 31, p=0.04, repeated measures ANOVA) D) Empirical cumulative distribution Function for the data shown in C. The number of spikes for the first half of the trace minus the spikes for the second half is shown for every cell.

## Discussion

The diversity of firing patterns upon step current injection that neurons present have been studied and catalogued for decades (McCormick et al., 1985; Markram et al., 2004; Ascoli et al., 2008). However, why neurons have evolved these diversity of responses remains still an open issue. As a form of rapid intrinsic plasticity which operates on the spike-timing, we shown here that CA3 neurons can switch between the main established suprathreshold discharge categories (Ascoli et al., 2008) after only a few minutes of subthreshold conditioning. This effect occurred upon input delivered either to their somata by activation of their synapses, or by direct intrasomatic current injection. The effect of conditioning was long-lasting, and abolished in the presence of PKA and PKC inhibitors, indicating that phosphorylation over the few minutes of conditioning is necessary for the changes in firing pattern. Hierarchical cluster analysis showed that the transitions observed are more likely towards adapting and intrinsic burst responses. Using a conductance-based neuron model and pharmacological blocking, we found that this shift can be explained by recruitment of calcium and M-type potassium conductances. The effect of the conditioning was also observed in CA3 neurons recorded from acute slices derived from juvenile mice, suggesting that this form of plasticity is a generic property of these cells. These results indicate that, on the time scale of minutes, CA3 neurons modulate their intrinsic firing pattern depending on their history of ongoing subthreshold activity.

### An intrinsic plasticity mechanism shapes the characteristic firing pattern of CA3 neurons

Neurons are plastic elements that can adjust their excitability by tuning membrane conductances in response to network activity (Turrigiano et al., 1994; Desai et al., 1999; Fan et al., 2005). This phenomena has been well characterized in the context of homeostatic plasticity (Turrigiano and Nelson, 2004). However, the time scale of those mechanisms typically extends over hours, and involves processes of gene expression (Lee et al., 2005), whereas in our experiments the changes were observed after only a few minutes of conditioning. Furthermore, we show that the effect is abolished by blocking phosphorylation, which also points towards a faster plasticity mechanism. More recent studies indicate that changes in intrinsic excitability occur on faster time scales (Aizenman and Linden, 2000; Paz et al., 2009; Mahon and Charpier, 2012; Brager and Johnston, 2007; Hyun et al., 2013). The variables considered are generally the firing rate or threshold of the cell, which are thought to facilitate synaptic hebbian learning (Titley et al., 2017). The neurons in our study significantly adapt their spiking dynamics, adding an extra timing dimension to the previous reports (in accordance to Grasselli et al. (2016) and Campanac et al. (2013)).

Most of the work on intrinsic plasticity require that the cell fire during the conditioning, whereas we observe that the firing patterns are modulated just by subthreshold input. Golowasch et al. (1999) already demonstrated that subthreshold current pulses could modulate the currents of lobster ganglion neurons, and so their discharge. However, this change required hours of stimulation. Brown and Randall (2009) and Sánchez-Aguilera et al. (2014) reported that subthreshold pulses or constant subthreshold depolarization increases the first ISI or reduces the overall excitability of the cell a few minutes after conditioning. In line with this work, Brown and Randall (2009) also reported that transient depolarizing pulses are more effective for the induction. Finally, other studies have shown that intrinsic cell properties can be affected by different neuromodulators (Brager and Johnston, 2007; Graves et al., 2012; Fujisawa et al., 2005). However, our changes were induced after direct subthreshold somatic conditioning, ruling out a synaptic cause.

The studies cited here were performed both in acute slices and in organotypic cultures. We find that the expression of the conditioning differed between these two preparations. Although induction of adaptation was found both cases, the bursting induction was qualitatively different, with acute neurons switching to delay burst responses (similar to Graves et al. (2012)) instead of intrinsic burst ones. These discrepancies may be given by a different channel distribution and kinetics, the species employed, the developmental time point or the activity levels of the two preparations (Kapoor et al., 1988; Moody and Bosma, 2005; Okamoto et al., 2014). As novel neurocentric forms of plasticity are being unraveled in behaving animals (Titley et al., 2017), further studies will be needed to determine whether the firing pattern plasticity is also found *in vivo* and if that is the case, which effect it may elicit on the complex dynamics of fully active circuits.

### Mechanisms of firing pattern transitions

Our modeling and pharmacological study suggests that the conductances supporting transitions through the firing pattern space of CA3 cells are *gKd*, *gKm* and *gCa* (coupled with *gCaK*). These candidates have previously been reported to shape the spiking response of hippocampal cells via activity dependent mechanisms. For example, *gCaT* is strongly associated with the switch to bursting mode upon status epilepticus (Kim et al., 2001; Su et al., 2002), while *gKd* is modulated by activity and influence the delay firing of the cell (Cudmore et al., 2010; Hyun et al., 2013; Saviane et al., 2003). Modulation of the M-type current upon activity has also been shown in CA1 (Wu et al., 2008) and in CA3 (Brown and Randall, 2009). Furthermore, rapid up- or down-regulation of ion channel maximal conductances via phosphorylation or vesicle modulation due to calcium signaling has been demonstrated extensively (Flavell and Greenberg, 2008; Davis et al., 2001; Zhang and Linden, 2003) and it has also been shown that the channels possess a complex of scaffold proteins containing protein kinases that could selectively regulate their conductance through phosphorylation (Davis et al., 2001). The exact rules that link the subthreshold input to the recruitment of the different conductances remain to be elucidated, but could likely depend on calcium dynamics and possibly the activation properties of the involved channels (Turrigiano and Nelson, 2004; Stemmler and Koch, 1999; Li et al., 2016).

One of the typical transitions that we observe is the switch towards bursting behaviors. We emphasize that this is not the only transition induced, but rather that special attention should be given to this bursting mechanism. It is known that some neurons present this dual behavior. For example, relay cells on the thalamus become bursty upon hyperpolarization because of T-type conductance inactivation (Sherman, 2001). In our case, the cells depolarized 5 mV in average, while kinase inhibitors blocked the effect, ruling out this hyperpolarizing cause.

### Functional implications of firing pattern modulation

Similar firing patterns are found in multiple species within the animal phyla (McCormick et al., 1985; Turrigiano et al., 1994; Yao and Wu, 2001), suggesting that they have a fundamental role in network computation. Shin et al. (1999) proposed that neurons that can dynamically adapt their output firing in response to their input statistics would have important advantages. By adjusting its threshold and dynamic range upon activity, a neuron could respond to stimuli over a broad range of amplitudes and frequencies without compromising sensitivity, maximizing the mutual information between its input and output discharge (Stemmler and Koch, 1999). Spike frequency accommodation has the characteristics of a high-pass filter (Benda and Herz, 2003). Since our conditioning stimuli occurred at constant frequencies, cells may have recruited a specific set of conductances that shift their integration properties to gain sensitivity in the new frequency range. Differences in filtering properties of brain stem neurons have also been shown to facilitate the extraction of spatial information from natural sounds (Remme et al., 2014) and most of the conductances that we identify in this study have reported to be frequency resonance candidates (Hutcheon and Yarom, 2000; Hu et al., 2002; Schreiber et al., 2004). These resonance properties of cells may also have important functional implications for neural activity and brain rhythms (Llinás, 1988; Buzsáki and Draguhn, 2004). When adjusting their discharge to more adapting patterns neurons may be changing mode from integrator to a coincidence detector (Prescott et al., 2008), helping not only to detect synchrony but also to transmit it to the network (Cudmore et al., 2010). Additionally, this fast plasticity of the firings may also be important for specific memory acquisition on the hippocampus (Kumaran et al., 2016; Benna and Fusi, 2016). CA3 is thought to be the generator of sharp wave ripples (SPW-R), a state where neurons cooperatively switch to presumably transfer memories to cortex (Buzsáki, 2015; Kumaran et al., 2016; Hunt et al., 2018). The stimulation protocol -resembling a SPW-R- may have push CA3 neurons to move to a different network state, similarly to Fujisawa et al. (2005), by possibly sensing the inputs and changing their spiking properties based on intrinsic plasticity rules (Srikanth and Narayanan, 2015).

Further studies will be needed in order to unravel the role that such firing pattern transitions may have for computations in neural circuits. A first step towards this goal must be to explore more generally how the form and frequency spectrum of somatic input signals on the long time scale affect the distinct firing patterns that neurons exhibit on the short scale. It appears that, after decades of study and cataloguing these patterns, the mystery is not whether they conform to classes or to a continuum but rather, what internal rules and computational advantages are that result neurons to converge onto these different discharge states.

### Grants

This work was supported by EU SECO grant EU216593 and ETH grant 2-73246-8 to Kevan A. C. Martin, Swiss National Science Foundation grant 31003A-143373 / 1 to U. Gerber and UZH Forschungskredit grant FK-18-119 to S. Soldado-Magraner.

### Disclosures

The authors declare no competing financial interests.

## Author Contributions

Conceptualization, S.S, F.B. and R.D.; Methodology, S.S. and F.B.; Software, S.S. and S.H.; Validation, S.S. and F.B.; Formal Analysis, S.S. and M.F.; Investigation, S.S., F.B. and S.H.; Resources, U.G. and K.A.C.M.; Data Curation, S.S.; Writing-Original Draft, S.S., M.F and R.D.; Visualization, F.B. and S.S.; Supervision, R.D. and M.F.; Project Administration, R.D.; Funding Acquisition, K.A.C.M., U.G. and S.S.

## Acknowledgements

We thank Kevan Martin and Dean Buonomano for critical comments on the manuscript, Beat Gähwiler for useful discussions, Ladan Egolf and Fritjof Helmchen for the help with the licences, Dubravka Göckeritz-Dujmovic for all the technical assistance, and Gabriela Michel and Marion Betizeau for proofreading the manuscript.

## Materials and Methods

All experiments were conducted in accordance with the guidelines and regulations of the Cantonal Veterinary Office of Zurich; License Nr 81/2014, 89/2013, 70/2016 and 156/2017.

### Electrophysiological Recordings

Rat hippocampal organotypic cultures (Gähwiler, 1981) of average postnatal age 21 days were transferred to a recording chamber and mounted on an upright microscope (Axioskop FS1; Zeiss). The cultures were superfused with an external solution (pH 7.4) containing (in *mM*) 148.8 *Na*^+^, 2.7 *K*^+^, 149.2 *Cl^−^*, 2.8 *Ca*^2+^, 2.0 *Mg*^2+^, 11.6 *HCO^−^*_3_, 0.4 *H*_2_*PO^−^*_4_, 5.6 D-glucose, and 10 *mg/l* Phenol Red. All experiments were performed at 34 *^◦^*C. Whole-cell recordings of CA3 neurons were obtained with patch pipettes (4-7 *M*Ω). Pipettes were filled (in *mM*) with 126 *K*-gluconate, 4 *NaCl*, 1 *MgSO*_4_, 0.1 *BAPTA − f ree*, 0.05 *BAPTA −Ca*^2+^, 15 glucose, 3 *ATP*, 5 *HEPES* (*pH* was adjusted to 7.2 with *KOH*) 0.1 *GTP*, and 10.4 byocitin. For acute experiments, 300 *µm* sagital hippocampal slices were prepared from mice ranging 15 to 22 days old. After decapitation under isoflurane anesthesia, the brain was removed and placed in oxygenated (95% *O*_2_, 5% *CO*_2_) and ice-cold high-sucrose artificial cerebrospinal fluid (ACSF) (pH 7.4) containing (in *mM*) 75 sucrose, 87 *NaCl*, 2.5 *KCl*, 7 *MgSO*_4_, 0.5 *CaCl*_2_ 26 *NaHCO*_3_, 1 *NaH*_2_*PO*_4_, and 10 D-glucose. Slices were cut in this solution and then transferred for recovery to oxigenated ACSF (*pH*7.4) at 37 *^◦^*C containing (in *mM*) 119 *NaCl*, 2.5 *KCl*, 1.3 *MgSO*_4_, 2.5 *CaCl*_2_ 26 *NaHCO*_3_, 1.25 *NaH*_2_*PO*_4_, and 10 D-glucose. After 30 min of recovery, slices were kept at room temperature for 1 hour before recording. During recording, slices were superfused with oxigenated ACSF.

The recording pipettes were manually positioned under microscope control. Recorded neurons were located mostly in the pyramidal cell layer. Electrophysiology and subsequent histology in a subset of the cells recorded suggest that the neurons described below include mostly pyramidal cells but also a subset of smooth cells.

Current-voltage relationships were determined by step command potentials and had duration of 1 second to ensure steady-state responses. Data were recorded using an Axopatch 200B amplifier (Molecular Devices). Series resistance was monitored regularly, and was typically between 5 and 15 *M*Ω. Cells were excluded from further analysis if this value changed by more than 20% during the recording. Junction potential and bridge was not corrected.

Mossy fibers were stimulated with a bipolar tungsten electrode. The intensity of the stimulus was constantly adjusted to evoke subthreshold post-synaptic potential responses of 15 mV on average in the recorded neuron.

Action potential discharges were evoked by injected current steps (−0.08 up to 1.8 nA; step increment 0.05 - 0.15 nA, depending on the input resistance of the recorded cell) each lasting from 3 to 5 seconds. After this control, the neurons were conditioned by mossy fibers activation, consisting of a double pulse (0.1 ms duration pulses, interval 10 - 20 ms) at a frequency of 1 Hz, repeated 500 times. Thus, the conditioning period was approximately 8 minutes. In a subset of experiments an interval of 60 ms was used. Immediately after this conditioning, the firing pattern of the neuron was assessed again using the same step protocol. The firing pattern of these cells was assessed every 10 minutes until 40 minutes of recording were completed to assess long-term effects of the plasticity. In a another subset of experiments, mossy fiber subthreshold responses were mimicked by injecting somatically and at a frequency of 1 Hz double step current pulses of 50 ms of duration and 20 ms of interstep interval. The amplitude of the pulse was adjusted in order to get a depolarization of 15 mV on average.

For any set of experiments, 1 cell per slice was recorded and 6 slices per rat on average (of either sex) were used for preparation of the organotypic cultures. A minimum of 8 cells were used for each set of experiments.

### Pharmacology

All drugs were applied to the slices via the perfusion system. Calcium currents were blocked by applying *NiCl*_2_ at 200 *µM*. D-type currents (Kv1 channels) were blocked by application of low concentrations (30 *µM*) of 4-aminopyridine (4 *−AP*). The M-current was blocked by application of *XE*991 at 10 *µM*, which targets *Kv*7*/KCNQ* channels (Brown and Randall, 2009). 4 *−AP*, *NiCl*_2_ and *XE*991 were purchased from Sigma-Aldrich (A78403, 339350, X2254). Times indicated in the figures reffer to the time elapsed since starting of the drug perfusion. The timing for the drug to reach the bath was estimated of 3-4 minutes.

### Histology

Hippocampal slice cultures were prepared for morphological assessment by fixing in freshly prepared 4% paraformaldehyde in 0.1 *M* phosphate buffer (*PB*) at *pH* 7.4 overnight at 4 *^◦^*C; washing three times in phosphate-buffered saline (*PBS*, 1.5 *mM KH*_2_*PO*_4_, 8.5 *mM Na*_2_*HPO*_4_, 137 *mM NaCl*, and 3 *mM KCl*, *pH* 7.4); and permeabilizing at room temperature in *PBS* that contained 10% heat-inactivated donkey serum, and 1% Triton X-100. Then they were incubated overnight at 4 *^◦^*C with streptavidin conjugated with Alexa (546*λ*). The cultures were washed again three times in *PBS*, and then mounted in Fluorostab (Bio-Science Products AG, Emmenbrucke, Switzerland) and coverslipped. High-resolution images were obtained using laser scanning confocal microscopy (Leica TCS SP2, Leica Microsystems, Heidelberg, Germany).

### Data analysis

Signals were digitized at 10 kHz and analysed off-line using pCLAMP 10 (Molecular Devices) and Matlab (MathWorks). Analysis of the voltage traces was performed similar to Chen et al. (2015). The average resting membrane potential of each neuron was estimated as the mean membrane potential during the first 100 ms of current-injection protocol (before injection of the step-current pulses). Input resistance was obtained by measuring the voltage drop across the hyperpolarizing trace of the step-current pulses. APs were located using median filtering, and the threshold was inferred as the point at which the derivative of the voltage trace exceeded 5 mV/ms. AP amplitude was measured from threshold-to-peak and AP afterhyperpolarization (AHP) from the threshold to through. Half-width was estimated as the full width at half-maximal amplitude. The fraction of spikes of a cell at a given time was computed by calculating the mean of the fraction of spikes of the individual current injections. At the population level the mean of this quantity was calculated.

Data was tested for normality using a one-sample Kolmogorov-Smirnov test. A paired t-test was used for statistical comparisons between conditions with normally distributed data and a two-sided Wilcoxon signed rank test was used otherwise. The exact p-values of the tests and the sample number for each experiment are indicated on the figure legends in the results section. A clustering analysis was also performed for the synaptic conditioning group, and it is described on the following section. All the statistical analysis were performed using Matlab (MathWorks).

### Cluster analysis of discharge traces

The firing patterns of the neurons were categorized by hierarchical clustering of their discharge patterns. The dataset consisted of all voltage traces recorded from neurons in response to step-wise current injections with different amplitudes, including recordings before and after conditioning. For any one neuron, the collection of responses to different current injections represents the signature of the electrical type. However, for inherent verification of our cluster procedure, we chose to treat each response independently. In this way successful clustering could be confirmed by its ability to assign responses from the same neuron into the same category.

The clustering measured similarity of a feature vector derived from the voltage traces. First the recorded voltage traces were converted into a time series of the instantaneous firing rates. The instantaneous firing rate at each spike was taken as 1/Inter-spike-Interval (ISI). Then the instantaneous rates where linearly interpolated across the spike times at 1 ms time intervals over 6 seconds (5 second current injection step, plus 1 second on and offset), and normalized by the maximum firing rate. Finally, a characteristic feature vector of a common length of 600 elements was obtained by down-sampling the interpolated rate traces by a factor of 10, in order to make them computationally tractable to the similarity measurement.

Similarity distances between pairs of traces were calculated using the Dynamic Time Warping (DTW) measure (Berndt and Clifford, 1994). DTW takes into account that two similar signals can be out of phase temporarily, and aligns them in a non-linear manner through dynamic programming (Keogh and Ratanamahatana, 2005). The algorithm takes two time series *Q* = 〈*q*_1_, *q*_2_,…, *q_n_*〉 and *C* = 〈*c*_1_, *c*_2_,…, *c_m_*〉 and computes the best match between the sequences by finding the path of indices that minimizes the total cumulative distance

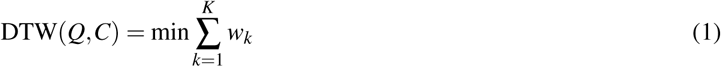

where *w_k_* is the cost of alignment associated with the *k^th^* element of a warping path *W*. A warping path starts at *q*_1_ and *c*_1_ respectively, and finds a monotonically increasing sequence of indices *i^k^* and *j^k^*, such that all elements *q_i_* in *Q* and *c _j_* in *C* are visited at least once, and for the final step of the path *i^end^* = *n* and *j^end^* = *m* holds. The optimal DTW distance is the cumulative distances *y*(*i, j*), corresponding to the costs of the optimal warping path *(q*_1_*,…, q_i_)* and *(c*_1_*,…, c _j_)*. This distance can be computed iteratively by dynamic programming:

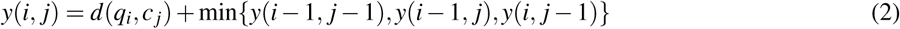

where *d*(*q_i_, c _j_*) is the absolute difference between the elements of the sequence. The optimal warping path is obtained by backtracking from the final element *y*(*n, m*), and finding which of the three options (increasing *i* only, increasing *j* only, or increasing *i* and *j* simultaneously) led to the optimal warping distance, until *i* = 1*, j* = 1 is reached. A warping window constraint of 10% of the vector size was chosen (Keogh and Ratanamahatana, 2005).

The pairwise DTW distances were used to perform hierarchical clustering by Ward’s algorithm (Ward Jr, 1963). The number of classes increases with the level of the hierarchy. We choose to cut the tree at a level that provided sufficient structure to interpret the hierarchy in terms of recognized response types (Ascoli et al., 2008).

Every recording for a given cell was treated as an independent observation, and could in principle be assigned to any cluster. If the electrophysiological state of the cell is expressed in all of its responses, then we expect that all the independent observations derived from that cell should be assigned to the same cluster. However, traces derived from current injections to the same cell in different conditions (pre- or post-stimulation) are expected to be assigned to different clusters if there is significant change in the underlying electrophysiological state.

In fact the independent traces did not cluster perfectly. Instead, the majority of independent observations derived from a given state clustered together and there were a few that fell into other clusters. Therefore, we chose to label the electrical type of each cell according to the cluster that contained the mode of the traces for one set of current injections. Cells for which no clear dominant cluster could be identified, e.g. because half of the traces fell into one cluster, and half of them into another, were labeled as unclassified. A cluster transition was recognized whenever the cell was assigned to different clusters before and after the stimulation protocol.

The analysis was performed using custom-written software in MatlabR2011b. The implementation of the DTW algorithm was obtained from Matlab Central http://www.mathworks.com/matlabcentral/fileexchange/43156-dynamic-time-warping-dtw.

### Neuron simulation model

A single cylindrical compartment, conductance-based neuronal model was used for all simulations. The length and diameter of the cylinder are set at equal dimensions to avoid spatial discretization problems in a single compartment (Cooley and Dodge, 1966; De Schutter and Bower, 1994). The passive properties associated with the model were obtained from Hemond et al. (2008). We set the length and diameter of our compartment to 50 *µ*m. The active properties were modeled by including appropriate voltage and calcium gated ion channels whose density and kinetics were obtained from experimental recordings performed in CA3 neurons (Hemond et al., 2008). All the conductances included in the model where obtained from this work, except for *gKd_slow_*, which had to be added in order to match the accelerating traces. We found that a 10 fold increase in the time constant of inactivation of *gKd* significantly improved the accelerating index. A similar slow *gKd* current matching these kinetics has actually been found in CA3 neurons (Luthi et al., 1996). The faster *Kd* current, has been previously reported both in cortex and hippocampus (Storm, 1988; Cudmore et al., 2010; Hyun et al., 2013; Miller et al., 2008). Throughout the manuscript, we refer to the channels as conductances. The simulations were performed using NEURON (Hines and Carnevale, 1997). We choose an integration step of 25 *µ*s, which was approximately 1% of the shortest time constant in the model. The voltage- and time-dependent currents followed the Hodgkin and Huxley formalism (1952):

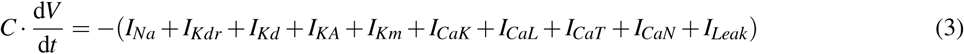

Each current *I_x_* is described by the equation

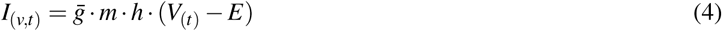

where 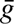 is the maximal conductance, *m* and *h* are activation and inactivation terms, *V* is the membrane potential, and *E* the reversal potential of the channel. The reversal potentials for *Na*^+^ and *K*^+^ were *E_Na_* = 50 mV and *E_K_*= -85 mV, respectively. The equations describing the different channel kinetics (*m, h*) for every current were obtained from Hemond et al. (2008). Following this reference, the three calcium conductances (T, N and L) were incorporated into a single parameter *gCa*.

The intracellular calcium dynamics were modeled as originally described by Hemond et al. (2008):

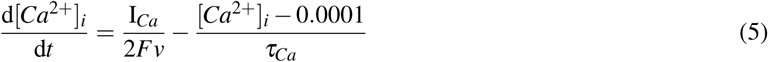

The first term of the above equation describes the change caused by *Ca*^2+^ influx into a compartment with volume *v*. F is the Faraday constant, *I_Ca_* is the calcium current and *τ_Ca_* is the time constant of *Ca*^2+^ diffusion.

The occasional decrease in spike amplitude seen in some of the experimental traces is probably due to sodium inactivation. We choose not to include this feature in the model, because it does not affect the overall dynamics of the spike discharge itself.

The model is available online at ModelDB https://senselab.med.yale.edu/ModelDB/enterCode.cshtml?model=228599

### Model Database of Traces

In order to get a conductance estimate for every voltage trace, we used the DTW algorithm to find the best fit to a database of voltage traces generated by the model. We varied the maximal conductances of our model into ranges that would contain our observed set of experimental voltage traces. The spiking conductances were left constant and *gCa*, *gCaK*, *gKm*, *gKd* and *gKd_slow_*were varied. The search of a valid conductance space was done manually, with the starting values provided by the report of Hemond et al. (2008) to reproduce CA3 firings. For the values and ranges used to generate the database see Table 1. A total of 100’000 conductance vectors were generated by combining the different conductances. The firing pattern of every conductance vector was produced at 8 different levels of step-current injection, obtaining a total of 800’000 voltage traces. An integration step of 0.2 ms was used. After generating the database non-spiking traces were removed, together with traces with saturating spikes. This led to a total of 73’024 voltage traces composing the conductance database. Every experimental trace, for both the control and conditioned case, was compared pairwise using the DTW algorithm with the set of voltage traces from the database. The 10 best fits were then selected in order to have an estimate of the conductance composition of the experimental trace.

The firing pattern of the model traces was simulated using 1 second of step current duration. Note that this differs from the time scale of our experimental traces, which were unraveled at 3-5 seconds of step current. Although generation of the traces for this longer duration was possible, the resulting firing patterns did not reproduce faithfully all the spiking dynamics encountered in the experiments. A change in channel kinetics (Hemond et al., 2008), an additional conductance, or a dendritic load could possibly solve the issue. The objective however was to gain an intuition on the possible conductance distribution changes induced by the conditioning. This together with computational reasons to generate the database led us to proceed with the simulations using a 1 second current step.

## Supplementary Material

**Figure S1.**
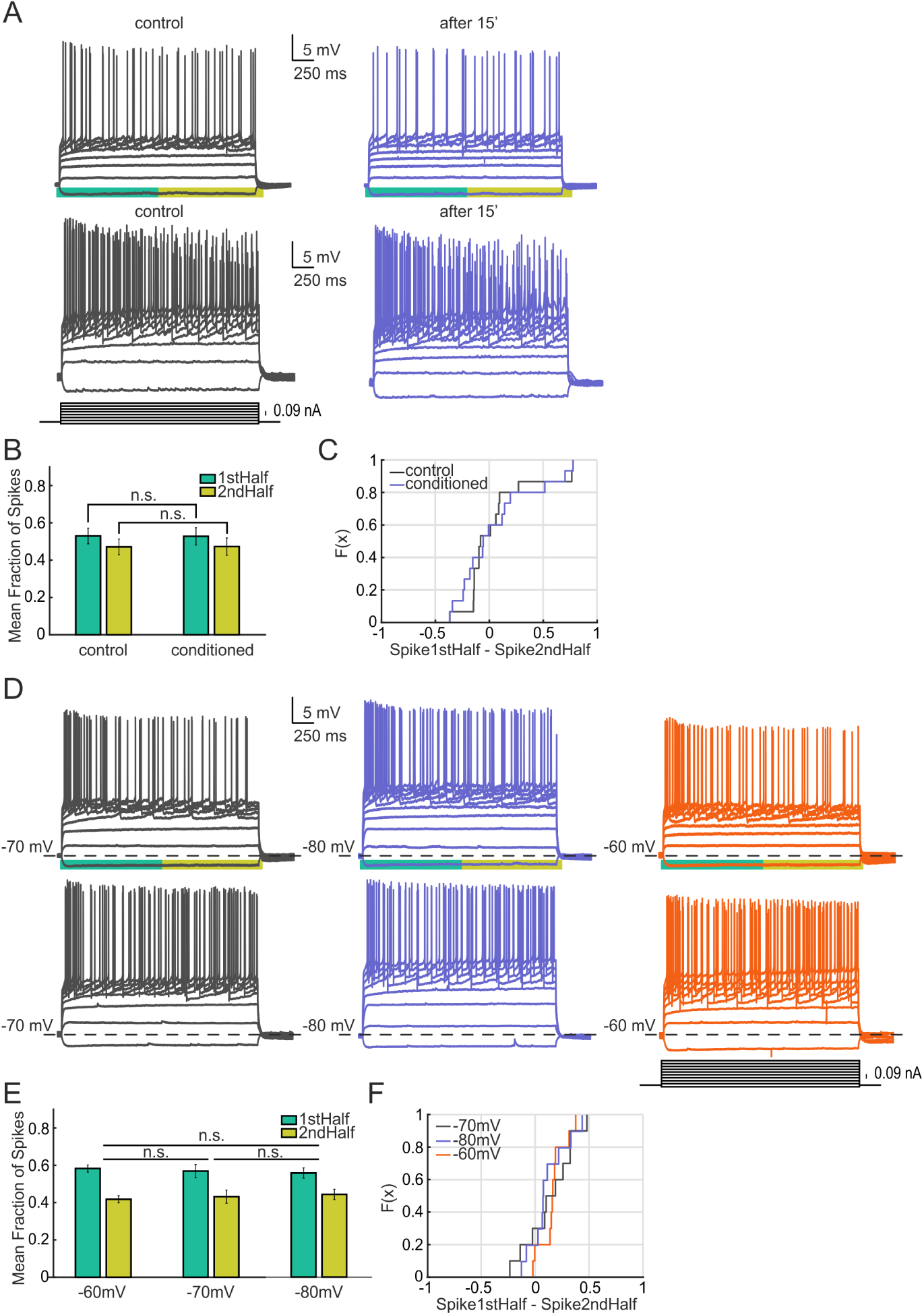
CA3 firing patterns are stable over time as well as to changes in resting membrane potential. Firing pattern transitions are not elicited by step current injection alone. A) Examples of two cells whose firing pattern have been measured by step-wise current injection (protocol showed in the inset). The cells do not show changes in firing pattern after 15 min of recording. B) Mean fraction of spikes for the population in the first and second half of the voltage trace for both control and conditioned cases. No significant redistribution on the fraction of spikes is observed (n = 15, p=0.583, two-sided Wilcoxon signed rank test). C) Empirical cumulative distribution Function for the data shown in B. Every individual case is represented as the number of spikes for the first half of the trace minus the spikes for the second half. D) Firing pattern transitions are not elicited by sustained shifts in membrane potential. Examples of two cells that have been hold at different membrane potentials through steady current injection (−70, −80 and −60 approximately). After changing the holding potential of the recorded neuron the firing pattern was measured by step-wise current injection (protocol showed in the inset). No transitions of firing pattern were observed at any of the different holding potentials. E) Mean fraction of spikes for the population in the first and second half of the voltage trace for every condition. No significant redistribution on the fraction of spikes is observed (Vm 60 vs 70, p=0.652; Vm 60 vs 80, p=0.084; Vm 70 vs 80, p=0.695) (n = 10, two-sided Wilcoxon signed rank test)). F) Empirical cumulative distribution function for the data shown in E. Every individual case is represented as the number of spikes for the first half of the trace minus the spikes for the second half.

**Figure S2.**
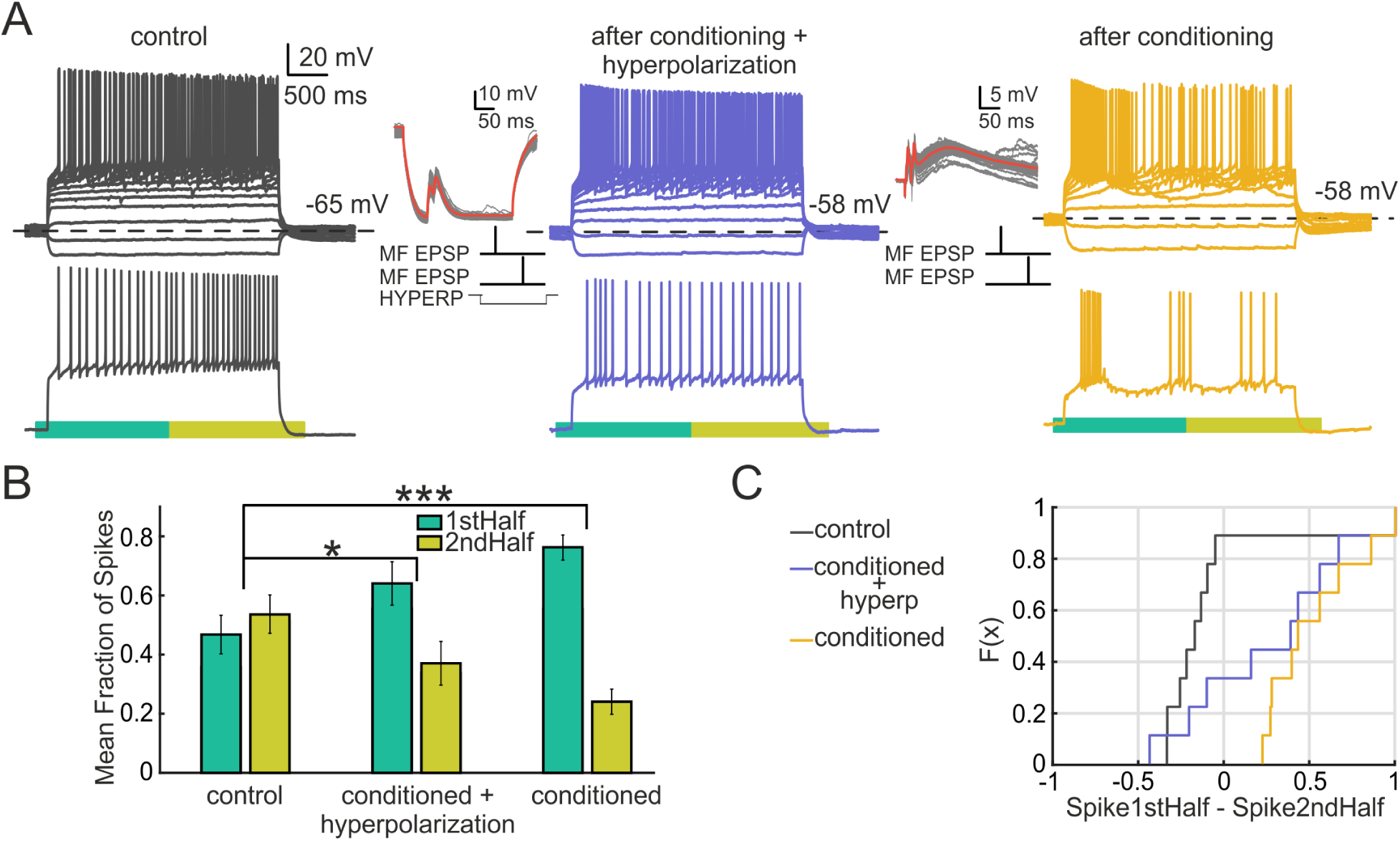
The effect of mossy fiber conditioning does not require somatic depolarization. Conditioning was performed while hyperpolarizing the cell with a negative current pulse. A) Cell that presents an accelerating pattern in control conditions (grey). Conditioning (MF double pulse, delta 20 ms given at 1 Hz) was elicited under the presence of an hyperpolarizing current step (protocol is shown in the inset, red line indicates the median). The cell changes its firing pattern to an non-adapting burst response. Thereafter, the cell is reconditioned via only the mossy fiber double pulse. After this conditioning the cell presents an intrinsic burst pattern. B) Mean fraction of spikes for the population in the first and second half of the voltage trace during the successive conditionings. A significant redistribution on the fraction of spikes is observed after the conditioning under the hyperpolarizing pulse (n=9, p=0.016, two-sided Wilcoxon signed rank test). After a second conditioning, without hyperpolarization, a stronger effect is found (n=9, p=0.008, two-sided Wilcoxon signed rank test). C) Empirical cumulative distribution Function for the data shown in B. The number of spikes for the first half of the trace minus the spikes for the second half is shown for every cell (n=9).

**Figure S3.**
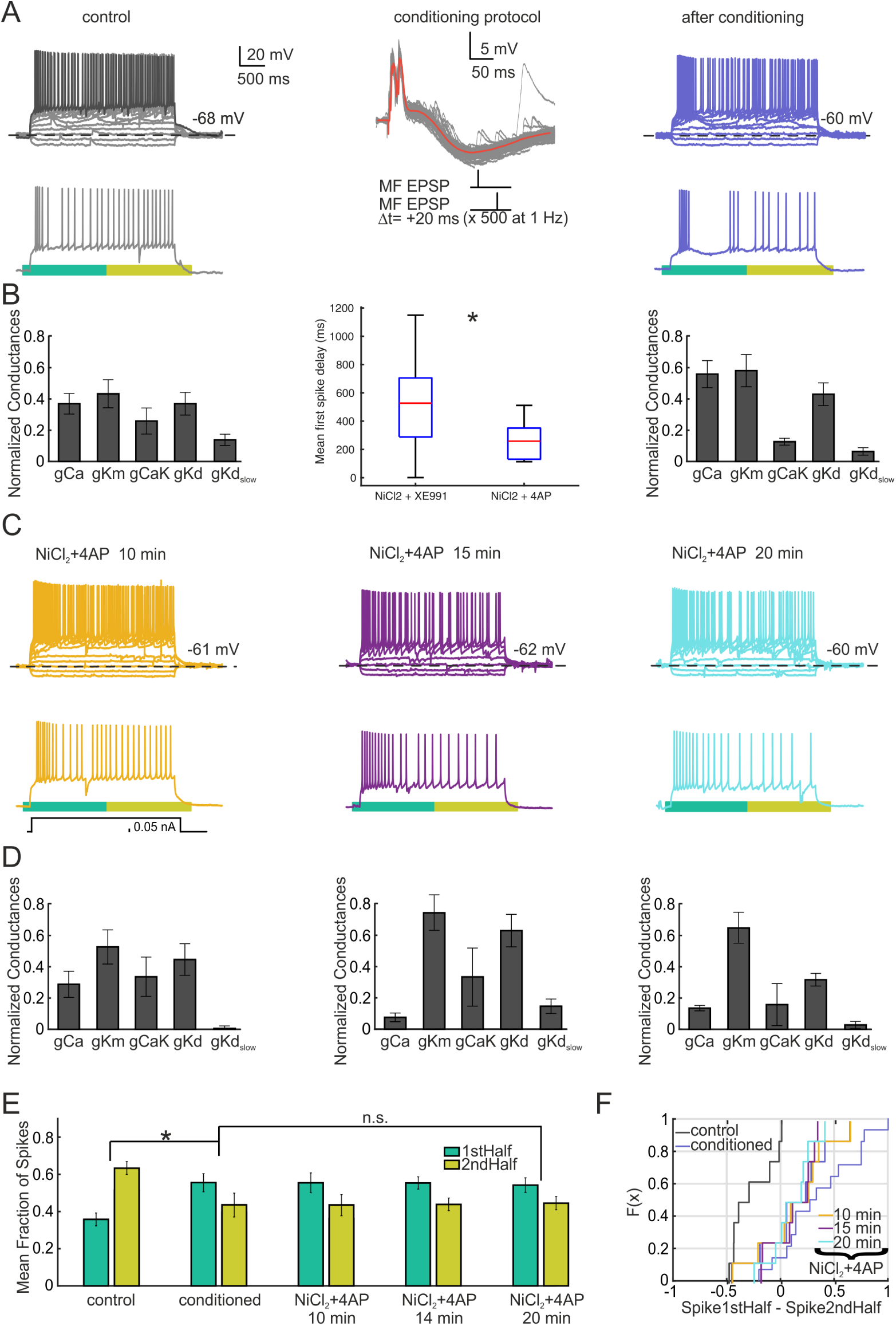
Inhibition of *Kv*1 and calcium channels reduces the delay of the traces but does not abolish the effect on the fraction of spikes. A) Cell with an adapting burst firing pattern in control conditions (grey) that switches to intrinsic burst after conditioning (blue). C) After 15 min of bath application of 4-AP and *NiCl*_2_ (at 30 and 200 *µM*) the cell switches to an adapting continuous pattern, with no delay (purple and blue traces). B) and D) Conductance model fits for every voltage trace. The middle panel shows the mean delay for the first spike when 4-AP is administrated with *NiCl*_2_ via the perfusion system, in comparison with *XE*991. A decrease on this delay by 4-AP is observed (n=8, p=0.02, two-sided Wilcoxon signed rank test). E) Mean fraction of spikes for the population in the first and second half of the voltage trace during the experiment. A significant redistribution on the fraction of spikes in favor of the first half is observed after conditioning (n=8, p=0.015, two-sided Wilcoxon signed rank test). Application of the drugs did not have an effect on the fraction of spikes (n = 8, p=0.98, repeated measures ANOVA) C) Empirical cumulative distribution function for the data shown in E.

